# Innocent until proven guilty: *Tannerella forsythia* may attenuate the virulence of *Porphyromonas gingivalis*

**DOI:** 10.1101/2025.09.05.674453

**Authors:** Mirosław Książek, Małgorzata Benedyk-Machaczka, Alicja Sochaj-Grzegorczyk, Danuta Mizgalska, Irena Waligórska, Magdalena Nowak, Anna Kaczor, Barbara Potempa, Andrew Fuchs, Ida B. Thøgersen, Jan J. Enghild, Carsten Scavenius, Piotr Mydel, Juhi Bagaitkar, Jan Potempa

**Author notes:** Contributed equally (Mirosław Książek, Małgorzata Benedyk-Machaczka). Jagiellonian Center of Innovation, Michala Bobrzynskiego 14 St., 30-348 Krakow, Poland (Alicja Sochaj-Grzegorczyk). Senior Corresponding Authors: Juhi Bagaitkar and Jan Potempa.

## Abstract

Periodontitis is caused by a dysbiotic microbiome beneath the gum line, primarily driven by the major pathobionts *Porphyromonas gingivalis* (*Pg*) and *Tannerella forsythia* (*Tf*). Their virulence depends on the excessive, uncontrolled activity of secreted proteases that sustain chronic inflammation, leading to the destruction of tissues supporting the tooth. Paradoxically, *Tf* also encodes multiple protease inhibitors, including miropin, a serpin with a broad range of targets. Here, we demonstrate that both native and recombinant miropin effectively inhibit lysine-specific gingipain (Kgp) and thiol protease (Tpr), impairing the growth of *Pg* in peptide-limited media and reducing its virulence *in vivo*. A rationally designed variant, RVK-miropin, also inhibited both lysine-specific and arginine-specific gingipains, fully suppressing *Pg* proliferation and virulence in a mouse infection model. Miropin is abundant on the surface of *Tf* and forms covalent inhibitory complexes with *Pg* proteases. In an oral gavage model of periodontitis, coinfection with wild-type *Tf* (but not a miropin-deficient mutant) and *Pg* significantly reduced alveolar bone loss caused by *Pg* alone. Miropin thus counteracts *Pg* virulence factors and host inflammatory responses, revealing an unexpected protective role for *Tf*. This challenges the traditional view of *Tf* as a primary periodontal pathogen, suggesting a context-dependent role as a microbial modulator within the dysbiotic biofilm. Beyond periodontal disease, the unique ability of miropin to inhibit structurally diverse proteases makes it a promising candidate for the development of new therapies that restore proteolytic balance in the periodontium.

## INTRODUCTION

Proteolytic processing is necessary for many biological processes, but the hydrolysis of peptide bonds by enzymes known as proteases is an irreversible post-translational modification and is therefore subject to strict regulation. The activity of many proteases is constrained by their synthesis as inactive proenzymes (zymogens) that are activated in proteolytic cascades (Obaha and Novinec, 2023; Kantyka et al., 2010; Song *et al*., 2022). In eukaryotes, but only rarely in prokaryotes, proteolysis is also regulated or limited by proteinaceous peptidase inhibitors, which account for 10% of the proteins in human serum and are plentiful in cells and tissues (Travis and Salvesen, 1983). An imbalance between proteases and their inhibitors underpins many pathological conditions, including periodontitis.

Periodontitis is an infection-driven chronic inflammatory disease characterized by excessive and uncontrolled proteolytic activity (Banbula *et al.*, 2000; Uito *et al.*, 20023; Radzki *et al.*, 2024). Tissue destruction in periodontitis results from a pathological inflammatory response to the dysbiotic microbial biofilm below the gum line. Dysbiosis reflects the synergistic interactions between periodontal bacterial pathobionts, where an antecedent colonizer creates conditions that favor subsequent colonization by other pathobionts, increasing the collective virulence of the microbial community and subverting host immune defenses. The proliferation of *Porphyromonas gingivalis* in the subgingival biofilm and the resulting increase in its proteolytic activity are considered necessary for the initiation of oral dysbiosis (Lamont et al., 2018; 2023).

The diverse proteases produced by *P. gingivalis* include abundant cysteine proteases known as gingipains, which specifically cleave peptide bonds adjacent to arginine (RgpA and RgpB) or lysine (Kgp) residues (Potempa *et al.*, 2000). Importantly, gingipain activity is unaffected by host protease inhibitors, which are themselves substrates of these bacterial enzymes (Abrahamson *et al.*, 1997; Plaza *et al*., 2017; Song *et al.*, 2022). Secreted gingipains can therefore disrupt several tightly regulated host proteolytic cascades, such as coagulation, fibrinolysis, kinin generation, and the activation of matrix metalloproteases, also triggering protease-activated receptor signaling pathways that promote nutrient influx and further inflammation (Guo *et al.*, 2010; Fitzpatrick *et al.*, 2009). The resulting conditions favor the proliferation of *P. gingivalis* and inflammophilic cohabitants such as *Tannerella forsythia* within periodontal pockets (Sheets *et al.*, 2008). In their natural habitat, *P. gingivalis* and *T. forsythia* coexist in close proximity within subgingival biofilms and infected periodontal tissues (Rajakaruna *et al*., 2018) and are frequently co-isolated from clinical samples (Mineoka *et al*., 2008; Naginyte *et al*., 2019). However, the molecular interactions between these bacteria, and the resulting synergistic or antagonistic outcomes, are not well understood.

Like *P. gingivalis, T. forsythia* is classified as a periodontal pathogen because it produces destructive enzymes such as the KLIKK proteases and PrtH, and is presumed to increase the overall virulence of the dysbiotic periodontal biofilm (Ksiazek *et al*., 2015a). But *T. forsythia* also secretes several potent protease inhibitors that can counteract a broad spectrum of peptidases. The effects of these inhibitors on the host and microbial community are unclear (Sharma, 2010; Ksiazek *et al*., 2015b). Protease inhibitors are rare in prokaryotes (both *Bacteria* and *Archaea*), with most species lacking protease inhibitor genes, and are also uncommon in protozoa and fungi. *T. forsythia* therefore stands out by producing multiple protease inhibitors, including α_2_-macroglobulin, an entire family of potempins (inhibiting co-produced KLIKK proteases), and the recently described protease inhibitor miropin (Ksiazek *et al.*, 2015a, 2022).

Miropin belongs to the serpin superfamily of serine peptidase inhibitors, but differs from archetypal serpins such as α_1_-antitrypsin, α_1_-antichymotrypsin and antithrombin III (which target serine proteases of defined specificity) by inhibiting a broad range of bacterial (subtilisin), plant (papain), and mammalian host proteases, including trypsin, plasmin, neutrophil elastase (NE), and cathepsin G (CatG) (Ksiazek *et al*., 2015b; Sochaj-Gregorczyk *et al.*, 2020). This broad specificity is attributed to the presence of multiple active sites within the miropin reactive center loop (RCL), which differs from the single-site architecture of typical serpins, and to the remarkable plasticity of its protein core, which can accommodate additional β-strands of varying lengths (Goulas *et al.*, 2017).

Given the broad inhibitory spectrum of miropin, its abundance on the surface of *T. forsythia*, and its ability to form stable, covalent complexes with diverse proteases (Ksiazek *et al*., 2015b; Goulas *et al.*, 2017), we hypothesize that it may also inhibit proteases secreted by *P. gingivalis*, thereby modulating the latter’s growth and virulence. Accordingly, we evaluated the ability of wild-type *T. forsythia* (wt-*Tf*) and a miropin-null isogenic mutant (*Tf*Δ*mir*) to inhibit gingipains and influence *P. gingivalis* virulence *in vitro* and *in vivo*. We also tested the activity of recombinant miropin variants containing the native -Val-Lys-Thr-(VKT-Mir) sequence at the RCL as well as the engineered variants -Val-Ala-Thr-(VAT-Mir) and -Arg-Val-Lys-(RVL-Mir). Our data challenge the classification of *T. forsythia* as a major periodontal pathogen by providing evidence that it attenuates the pathogenicity of *P. gingivalis* in co-infection models.

## MATERIALS AND METHODS

### Enzymes, inhibitors, substrates, and activity titration

Unless otherwise stated, all reagents were purchased from Millipore-Sigma (St Louis, MO, USA). The proteases RgpB, Kgp and Tpr were purified from *P. gingivalis* as previously described (Veillard *et al.*, 2015; Staniec *et al.*, 2015; Sztukowska *et al.*, 2012). Human neutrophil elastase (NE) human α_2_-macroglobulin were obtained from Athens Research and Technology (Athens, GA, USA). We also used trypsin, and protease substrates Suc-LLVY-AMC, *N*-tosylo-GPK-pNA, Bz-Arg-pNA and Ac-Lys-pNA. RgpB, Kgp, and Tpr were active-site titrated with *N*-α-tosyl-L-lysine chloromethyl ketone (TLCK) hydrochloride. NE were titrated with α_2_-macroglobulin, assuming one native inhibitor molecule inhibits two protease molecules, using FITC-casein (Thermo Fisher Scientific, Waltham, MA, USA) as the substrate. The active site concentration of α_2_-macroglobulin was determined using trypsin titrated with 4-nitrophenyl 4-guanidinobenzoate. Before use, Kgp, RgpB and Tpr were activated in gingipain assay buffer (GAB) (50 mM Tris-HCl pH 7.5, 150 mM NaCl, 2.5 mM CaCl_2_, 0.02% NaN_3_) containing 10 mM L-cysteine (GAB-C) for 15 min at 37 °C.

### Stoichiometry of inhibition (SI) and association rate constant (*k_ass_*)

To determine the SI, proteases Kgp, RgpB, NE (all 10 nM), and Tpr (100 nM) were mixed with increasing concentrations of miropin in wells of a microtiter plate in 100 μL GAB-C (Kgp, RgpB and Tpr) or GAB with 0.05% Pluronic F-127 (GAB-P). After pre-incubation at 37 °C for 10 min, we added 100 μL of substrate: 150 μM *N*-tosylo-GPK-pNA (Kgp), 25 μM Suc-LLVY-AMC (Trp), 200 μM Bz-Arg-pNA (RgpB), 1 mM MeSuc-AAPV-pNA (NE), or 1 mM Suc-AAPF-pNA (CatG). The residual protease activity for all except Trp was measured at 410 nm using a Sunrise microplate reader (Tecan, Männedorf, Switzerland). Tpr activity was measured by fluorimetry (excitation = 355 nm, emission = 460 nm) using a SpectraMax Gemini EM microplate reader (Molecular Devices, San Jose, CA, USA). To determine the *k_ass_*, constant substrate concentrations and increasing concentrations of miropin in a total volume of 100 μL in microtiter plate wells were mixed with 100 μL of protease solutions (2 nM Kgp or RgpB, 20 nM Tpr), and substrate hydrolysis was monitored as described above using the same buffers. Data were analyzed as previously described (Ksiazek, 2015b).

### Bacterial strains

Wild-type strains of *T. forsythia* ATCC 43037 (wt-*Tf*) and *P. gingivalis* W83 (wt-*Pg*) were purchased from the American Type Culture Collection (ATCC; Rockville, MD, USA). We also used a Kgp deletion mutant (*Pg*Δ*kgp*) (Sztukowska *et al.*, 2004) and a gingipain-null W83 strain (*Pg*ΔKΔRAB) (Rapala-Kozik *et al*., 2011), as well as an isogenic miropin-null mutant (*TfΔmir*) prepared specifically for this study.

### Bacterial growth and culture fractionation

The wt-*Pg*, *Pg*Δ*kgp,* and *Pg*ΔKΔRAB strains were cultured in enriched tryptic soy broth (eTSB; Millipore-Sigma) supplemented with 5 g/L yeast extract, 5 mg/L hemin, 0.5 g/L L-cysteine, and 2 mg/L menadione (*Pg* eTSB). The wt-*Tf* and *Tf*Δ*mir* strains were grown in eTSB supplemented with 5% horse serum (Gibco, Thermo Fisher Scientific), 0.25 g/L L-cysteine, 1 mg/L hemin, 0.5 mg/L menadione, and 10 mg/L *N*-acetylmuramic acid (*Tf* eTSB). Bacterial cultures were incubated in an anaerobic chamber (Don Whitley Scientific, Bingley, UK) in an atmosphere of 90% nitrogen, 5% carbon dioxide, and 5% hydrogen at 37 °C. The solid medium for *P. gingivalis* and *T. forsythia* was the appropriate eTSB plus 1.5% agar, supplemented with 5% defibrinated sheep blood for *P. gingivalis* (Graso Biotech, Starogard Gdanski, Poland). *Tf*Δ*mir* was grown in the presence of 5 μg/mL chloramphenicol, *Pg*Δ*kgp* in the presence of 1 μg/mL tetracycline, and *Pg*ΔKΔRAB in the presence of 1 μg/mL tetracycline, 5 μg/mL chloramphenicol, and 5 μg/mL erythromycin. All bacterial strains were cultured to log phase, which was OD_600_ = 0.4–0.6 for *T. forsythia* and OD_600_ = 0.8–1.2 for *P. gingivalis*. Whole bacterial cells were harvested by centrifugation (8000 x *g*,10 min, 4 °C for *P. gingivalis*; 5000 x *g*, 10 min, 4 °C for *T. forsythia*), washed twice in sterile phosphate-buffered saline (PBS), and resuspended in 10% of the initial culture volume of GAB-C. *T. forsythia* supernatant was passed through a 0.45-µm syringe filter followed by ultracentrifugation (100,000 x *g*, 1.5 h, 4 °C) to harvest outer membrane vesicles (OMVs). The OMVs were washed twice in GAB and resuspended in 1% of the initial culture volume of GAB-C. To obtain the cell envelope fraction containing the outer and inner membrane and peptidoglycans, *T. forsythia* whole cells were sonicated, followed by ultracentrifugation as above. The supernatants were discarded, and the pellet was resuspended in GAB-C.

### Generation of the miropin knockout strain of *T. forsythia* (*Tf*Δ*mir*)

A suicide plasmid designed to replace the *Miropin* coding sequence (CDS) with a chloramphenicol resistance cassette (chloramphenicol acetyltransferase gene, *cat*) was synthesized commercially. The plasmid contained a 728-bp fragment of the *T. forsythia* genome upstream of the *Miropin* start codon and a 786-bp fragment downstream of the *Miropin* stop codon, separated by the 660-bp CDS of the *cat* gene. These elements were transferred to pUC57 by integration at the multiple cloning site to produce pΔMiropin and introduced into electrocompetent *T. forsythia* cells. Positive clones were selected on eTSB agar plates with 10 µg/mL chloramphenicol and were sequenced for confirmation.

### Site-directed mutagenesis and purification of miropins

Miropin mutants with amino acid substitutions at the ^367^VKT^369^ sequence motif of the RCL were generated using plasmid pGex-6P-1 containing cloned miropin (Goulas *et al*., 2017), appropriate oligonucleotides, and the QuikChange Lightning site-directed mutagenesis kit (Stratagene, La Jolla, CA, USA). Primers VAT-mirF: 5’ CCGTAGAAATGGTAGCAACGTCATCCCCCTC 3’ and

VAT-mirR: 5’ GAGGGGGATGACGTTGCTACCATTTCTACGG 3’ for VAT-mir

VKT-mirF; 5’ CGTAGAAATGGTAGCAACGTCATCCCCCTC 3’ and

VKT-mirR: 5’ GAGGGGGATGACGTTGCTACCATTTCTACGG 3’ for VKT-mir

RVK-mirF: 5’ GCCGTAGAAATGCGCGTGAAATCATCCCCCTCTACC 3’ and

RVK-mirR: 5’ GGTAGAGGGGGATGATTTCACGCGCATTTCTACGGC 3’ for RVK-mir were synthesized by Genomed (Warsaw, Poland). Each construct was verified by DNA sequencing. Recombinant wild-type miropin (VKT-mir) and its variants (RVK-mir and VAT-mir) were expressed in *Escherichia coli*, then purified by affinity chromatography on glutathione-Sepharose columns and size exclusion chromatography on HiLoad 16/60 Superdex 75 pg columns as previously described (Goulas *et al.*, 2017).

### Miropin effect on *P. gingivalis* growth

*P. gingivalis* cells were harvested, washed twice in Dulbecco’s modified Eagle’s medium (DMEM; Gibco) containing 1% (w/v) bovine serum albumin (BSA), and suspended in DMEM containing 1% BSA as well as hemin, L-cysteine, and menadione at the same concentrations found in *Pg* eTSB. The cultures were pre-equilibrated in an anaerobic atmosphere and allowed reach OD_600_ = 0.4. Tubes were sealed in an anaerobic chamber and bacterial growth in the presence or absence of miropin was monitored by measuring the OD_600_ every hour for 24 h using a DiluPhotometer (Implen, München, Germany).

### Inhibition of gingipains on the *P. gingivalis* cell surface

*P. gingivalis* cells were washed and suspended in GAB-C with the OD_600_ adjusted to 1.0. We mixed 20 µL of this suspension with 100 µL GAB-C containing increasing concentrations of miropin in a microtiter plate. After 15 min at 37°C, the residual activity of Kgp, RgpB, and Tpr was recorded as described above for SI determination. Similarly, increasing amounts of *T. forsythia* whole cell suspension (OD_600_ = 1.0) or OMVs were pre-incubated with 1 nM Kgp in a microtiter plate (total volume 100 μL) for 15 min at 37 °C in GAB-C, and residual activity was determined using 1 mM Ac-Lys-pNA.

### Gel electrophoresis and immunoblotting

Samples mixed 1:1 (v:v) with 2× non-reducing or reducing sample buffer (plus 50 mg/mL dithiothreitol (DTT) for samples with chymotrypsin) were resolved by SDS–PAGE using 10% (T:C ratio, 33:1) gels and the Tris-Tricine buffer system (Schägger and von Jagow, 1987). For immunoblot analysis, proteins were transferred to 0.22-μm pore PVDF membranes (Bio-Rad, Hercules, CA, USA) using a semi-dry transfer unit (40 min at 15 V in 50 mM Tris, 40 mM glycine, 0.04% SDS, 10% methanol) and non-specific binding sites were blocked with 5% (w/v) skimmed milk in 20 mM Tris-HCl pH 7.5, 0.5 M NaCl, 0.1% Tween-20 (TTBS). Blots were probed at 4 °C overnight with a rabbit polyclonal anti-miropin antibody diluted 1:1000 in blocking solution (0.6 μg/mL). The antibody was generated by Kaneka Eurogentec (Seraing, Belgium) using recombinant miropin as the antigen. The blots were then washed and incubated for 1 h at room temperature with an HRP-biotin-conjugated goat anti-rabbit polyclonal secondary antibody (BioShop, Burlington, Canada) diluted 1:15,000 in blocking solution. Signals were developed using Pierce ECL Western Blotting Substrate (Thermo Fisher Scientific) before imaging.

### Detection of covalent inhibitory complexes

Miropin at concentrations of 4.8, 2.6, and 25 μM was incubated with increasing concentrations of proteases in 20 μL GAB-C for 15 min at 37 °C. We then added 2 μL of 2 mM solutions of TLCK, incubated for 5 min at room temperature, and resolved the samples by SDS-PAGE. Protein identities were determined by mass spectrometry. Diluted suspensions of *T. forsythia* whole cells or OMVs in GAB-C were mixed with *P. gingivalis* whole cells in 20 μL GAB-C and incubated for 15 min at 37 °C. Residual gingipain activity was inhibited by adding 2 μL of 5 mM TLCK for 5 min at room temperature, and samples were analyzed by immunoblotting with anti-miropin antibodies as described above.

### Determination of the cleavage sites in the miropin RCL

Miropin (30 or 40 μM) was mixed with Kgp (10 μM) or Tpr (7.5 μM) in a total volume of 20 μL GAB-C and incubated for 15 min at 37 °C. The mixtures were resolved on 18% polyacrylamide gels and transferred to PVDF membranes in 10 mM CAPS, 10% methanol (pH 11). The N-terminal sequences of stained peptides with a molecular weight of ∼4.5 kDa were obtained by automated Edman degradation using a Procise 494HT amino acid sequencer (Applied Biosystems, Carlsbad, CA, USA) as previously described (Ksiazek *et al.*, 2015b).

### Dot blots and flow cytometry

*P. gingivalis* and *T. forsythia* cells were harvested, washed, and resuspended in cold PBS containing 200 µM TLCK and 1×cOmplete, Mini, EDTA-free Protease Inhibitor Cocktail at an OD_600_ of 1.0. Half of each intact cell suspension was supplemented with 0.05% SDS (*P. gingivalis* only) and sonicated to disrupt the cells. We then spotted 5 μL of the intact or sonicated cell suspensions onto 0.22-μm nitrocellulose membranes (Bio-Rad), which were air-dried, probed with a rabbit polyclonal anti-miropin antibody in blocking solution (0.6 μg/mL) and developed using ECL as described above. For flow cytometry, the intact cell suspension was suspended in flow cytometry buffer or FCB (PBS with 1.5% BSA, 1 mM TLCK, and 1× cOmplete Protease Inhibitor Cocktail). We transferred 100 μL of the suspension in FCB to a 96-well conical-bottom plate and incubated for 30 min on ice with the polyclonal rabbit anti-miropin antibody (30 μg/mL). Bacterial suspensions were centrifuged at 1000 x g and washed once with FCB before incubation with an Alexa Fluor 488-conjugated goat anti-rabbit secondary antibody (diluted 1:200) or only with Streptavidin−FITC (diluted 1:200). The cells were washed twice, resuspended in 250 μL FCB, and analyzed on a FACSCalibur flow cytometer with CellQuest software (BD Biosciences, Franklin Lakes, NJ, USA). Data were analyzed using FlowJo software (FlowJo, Ashland, OR, USA).

### Experimental animals and ethical approval

Specific pathogen-free (SPF) female BALB/c mice, 8–10 weeks of age, were purchased from Janvier Labs (Le Genest-Saint-Isle, France). Mice were housed in individually ventilated cages and fed a standard laboratory diet and water *ad libitum* under SPF conditions at the animal care facility, Faculty of Biochemistry, Biophysics, and Biotechnology, Jagiellonian University, Kraków, Poland. Mice were maintained at 22 ± 2 °C and 60 ± 5% relative humidity with a 12-h photoperiod. Control and infected mice were housed in separate cages. All animal experiments were reviewed and approved by the I Regional Ethics Committee on Animal Experimentation, Kraków, Poland (approval no: 131/2018).

### Subcutaneous chamber model

Mice were anesthetized by the intraperitoneal injection of ketamine (200 mg/kg VetaKetam; VetAgro, Lublin, Poland) and xylazine (10 mg/kg Sedasin; Biowet Puławy, Puławy, Poland), and the eyes were lubricated with Puralube Vet (Pharmaderm, Tunis, Tunisia). Following general anesthesia, using sterile instruments, the skin was incised on the right side of the back, and a sterile titanium-coil chamber was implanted subcutaneously. After 10 days, bacteria were injected into the lumen of the subcutaneous chamber in the following combinations: (*i*) 10^9^ CFUs of wt-*Pg* and 250 µg of VKT-miropin, RVK-miropin, or VAT-miropin, or (*ii*) 1 × 10^9^ CFUs wt-*Pg* or 1 × 10^9^ CFUs *Pg*Δ*kgp* or 1 × 10^9^ CFUs wt-*Tf* or 1 × 10^9^ CFUs *TfΔmir* or 1 × 10^9^ CFUs wt-*Pg* plus 1 × 10^9^ CFUs wt-*Tf* or 1 × 10^9^ CFUs wt-*Pg* with 1 × 10^9^ CFUs *Tf*Δ*mir* in a total volume of 100 µL. Control mice were sham-infected by injecting 100 µL of sterile PBS. Clinical signs of infection, morbidity, and mortality were assessed as indicators of *P. gingivalis* virulence over a 6-day period. The chamber fluid (exudate) was collected at different time points after inoculation using a sterile 1-mL syringe and resuspended in PBS (pH 7.4). Ten-fold serial dilutions were plated on blood agar plates, cultured for 7–10 days in an anaerobic chamber at 37°C, before CFUs were determined. Chamber exudates were also assessed for gingipain activity and myeloperoxidase (MPO) activity.

### Gingipain activity in chamber fluids

To measure the proteolytic activity of gingipains, 2 µL of chamber fluid was added to 98 µL TNCT buffer (100 mM Tris-HCl pH 7.5, 150 mM NaCl, 5 mM CaCl_2_, 0.05% Tween-100), supplemented with 10 mM L-cysteine in a microtiter plate. Following a 5-min pre-incubation, we added substrate for Rgp (Bz-Arg-pNA) or Kgp (*N*-p-Tos-GPK-*p*NA). The resulting mixture contained 200 μM substrate, 5% DMSO, and 10 mM L-cysteine. The gingipain-mediated release of *p*-nitroaniline from the substrates was monitored as an increase in absorbance (405 nm) over time using a SpectraMax Gemini microplate reader.

### Myeloperoxidase (MPO) assay

Neutrophil influx into the chamber fluids was analyzed using MPO as a surrogate marker. Briefly, 2 µL of chamber fluid was mixed with dianisidine reagent (16.7 mg dianisidine, 90 mL distilled water, 10 mL potassium phosphate buffer, 50 µL 0.29 M hydrogen peroxide) and incubated for 5 min at room temperature. The reaction was terminated by adding 50 µL 2 M H_2_SO_4_. The change in absorbance over time was measured at 450 nm, and MPO activity was expressed as a percentage of the control value (PBS-treated animals).

### Oral gavage model

Mice were pre-treated with 870 μg/mL sulfamethoxazole and 170 μg/mL trimethoprim to deplete oral microflora for 8 days. On day 10, mice were orally infected with 1 × 10^10^ CFU wt-*Pg* or wt-*Tf* or wt-*Pg* plus wt-*Tf* or wt-*Pg* plus *TfΔmir* suspended in 2% sterile CMC by oral gavage on alternate days for a total of 2 weeks. Mice were euthanized 4 weeks after the last bacterial inoculation. Alveolar bone loss was determined by using micro-computed tomography (µCT) to measure the distance from the cementoenamel junction (CEJ) to the alveolar bone crest (ABC) (Abe and Hajishengallis, 2013).

### Statistical analysis

All statistical analyses were performed using GraphPad Prism version 10.4.1 (GraphPad Software, Boston, MA, USA). Outliers were identified using the ROUT method (Q = 0.2%) and excluded from subsequent analyses, except for the datasets presented in Fig. 4 (panels A and B) and Fig. 5 (panels A, C, and E), where no outliers were eliminated. Data are presented as mean ± standard deviation (SD), except for Fig. 5C and Fig. 5D, where values are shown as mean ± standard error of the mean (SEM). Survival curves were analysed using the Kaplan–Meier method, and statistical differences between groups were assessed using the log-rank (Mantel–Cox) test (Fig. 4A and Fig. 5A). Group comparisons were performed using Welch’s t-test or one-way ANOVA followed by Dunnett’s multiple comparisons test, as appropriate. A p-value ≤0.05 was considered statistically significant (\**p* < 0.05, **p < 0.01, \*\*\**p* < 0.001; ns = not significant).

## RESULTS

### Recombinant miropin inhibits *P. gingivalis* proteases Kgp and the Tpr

The protease inhibitor miropin, produced by *T. forsythia*, is unique in the serpin superfamily because it inhibits proteases with diverse specificities, catalytic mechanisms, and evolutionary origins (Ksiazek *et al.*, 2025, 2020; Goulas *et al.*, 2017). Therefore, we screened the inhibitory activity of recombinant wild-type miropin (VKT-mir) against cysteine endopeptidases secreted by *P. gingivalis.* At a 10-fold molar excess, miropin completely inhibited the proteolytic activity of purified Kgp and Tpr but not RgpB (**Fig. 1A**). Miropin inhibited Kgp and Tpr in a concentration-dependent manner, with SI values of 1.9 and 9.3, respectively (**Fig. 1 B,C**), indicating the number of molecules of the serpin consumed in the reaction to form a stable covalent inhibitory complex. The formation of such complexes was supported by SDS-PAGE and confirmed by peptide mass fingerprinting of protein bands designated as complexes (Suppl. **Fig. S1A**). Finally, N-terminal sequencing of the ∼5-kDa peptide released from miropin during complex formation and/or non-productive RCL cleavage revealed that Kgp attacked the Lys^368^-Thr (P2-P1) peptide bond in the RCL as expected by the exclusive specificity of this protease for Lys-Xaa peptide bonds (Pike *et al.*, 1994). In contrast, Tpr cleaved the RCL at two positions: Lys^368^-Thr and Ser^373^-Thr (P4′-P5′) (Suppl. **Fig. S1B**). The latter is too far downstream from the P2-P1 reactive site bond (RSB) to achieve covalent complex formation, corroborating the high SI. Therefore, the Tpr protease is inhibited by miropin at Lys^368^-Thr as the RSB.

**Figure 1.**
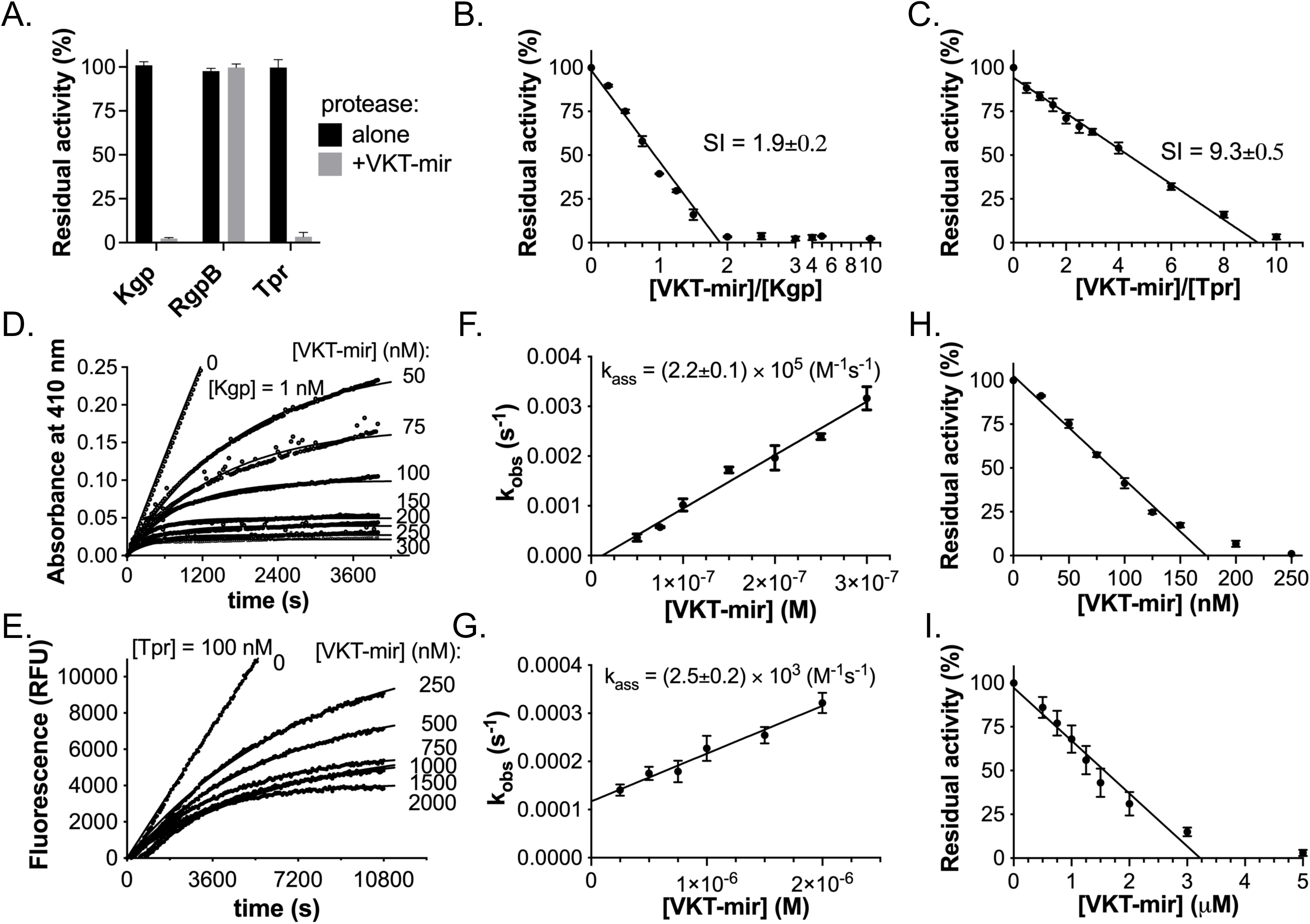
Miropin inhibits soluble and *P. gingivalis* surface-associated Kgp and the Tpr protease but not Rgp. Purified RgpB, Kgp and Tpr were incubated with 10-fold molar excess of miropin (**A**) or increasing concentrations of miropin for 15 min, and the residual activity of Kgp (**B**) and Tpr (**C**) was measured using specific substrates. The residual activity of each enzyme was plotted against the miropin/protease molar ratio to determine the stoichiometry of inhibition (SI). To determine the association rate constant (*k_ass_*), substrate hydrolysis by Kgp (**D**) and Tpr (**G**) in the presence of constant amounts of substrate and increasing concentrations of VKT-mir was recorded over time. We plotted *k_obs_* (observed associated rate constant) against inhibitor concentrations, and the *k_ass_* for Kgp (**E**) and Tpr (**H**) were computed from the slope of the linear curve fitted to the data points corrected for SI and K_M_. (**F**, **I**) A *P. gingivalis* cell suspension (OD_600_ = 1.0) was pre-incubated with increasing concentrations of miropin, and the residual activity of Kgp (**F**) and Tpr (**I**) was determined and plotted against the miropin concentration. Data are means + SD (n = 3).

Irreversible protease inhibitors presumably control proteolysis by reacting with their target protease with a *k_ass_* of at least 10^5^ M^-1^ s^-1^ (Bieth, 1984). Progress curve analysis revealed *k_ass_* values of 2.2 x 10^5^ and 2.5 x 10^3^ M^-1^s^-1^, respectively, for the inhibition of Kgp (**Fig. 1D,E**) and Tpr (**Fig. 1G,H**). The relatively slow *k_ass_* and high SI value argue that the inhibition of Tpr by miropin has no pathophysiological significance *in vivo* because it would take too long and consume too many molecules of miropin to inhibit one molecule of the protease. Nevertheless, the activity of both Kgp and Tpr on the surface of *P. gingivalis* (whole bacterial cells) is sensitive to inhibition by recombinant miropin, and the inhibitor could be used to titrate the level of these proteases on the bacterial cell surface (**Fig. 1F, I**).

These results confirm that *T. forsythia* miropin is an unusual serpin that inhibits not only serine proteases and papain (clan CA, family C1) (Goulas *et al*., 2017) but also different families and clans of cysteine peptidases. Tpr belongs to family C2 (clan CA) of peptidases and shares similarity with calcium-regulated calpains (Staniec *et al*., 2015, 2023), whereas the protease domain of Kgp is structurally distinct from papain and calpain, resembling caspases and belonging to cysteine peptidase family C25, clan CD (de Diego *et al.*, 2014). The highly efficient inhibition of the soluble and cell surface-associated forms of Kgp (**Fig. 1B,I**) suggests that the inhibitor can modulate the virulence of the subgingival microbiome, given that Kgp is an essential *P. gingivalis* virulence factor (O’Brien-Simpson *et al.*, 2000; Pathirana *et al*., 2006) and that *T. forsythia* and *P. gingivalis* coexist in pathological periodontal pockets and infected gingival tissue (Socransky *et al*., 1998; Rajakaruna *et al*., 2018).

### Miropin is exposed on the *T. forsythia* cell surface

We previously used SignalP and LipoP to confirm that the nascent miropin translation product is a lipoprotein (Ksiazek *et al.*, 2015b). In diderm bacteria, lipoproteins exported into the periplasm via the Sec system are anchored into the inner or outer membrane and face the lumen of the periplasm (Bos *et al.*, 2007). Alternatively, a lipoprotein can be translocated across the outer membrane and displayed on the bacterial surface, anchored into the membrane’s outer layer (May and Grabowicz, 2025; Cole *et al.*, 2021). The subcellular location of miropin cannot be predicted reliably, so we determined its cellular location by dot blot analysis of intact and sonicated wt-*Tf*, with sonicates of *P. gingivalis* and the *Tf*Δ*mir* mutant as controls. The intensity of the miropin signal was identical in the intact and sonicated cells (**Fig. 2A, left panel**). Notably, no signal was detected for *Tf*Δ*mir* and *P. gingivalis* cells regardless of sonication, confirming the antibody’s specificity for miropin. In *P. gingivalis,* the inner membrane protein MmdC is the only naturally biotinylated protein that can serve as an inner membrane marker (Lasica *et al.*, 2016). A biotin carboxyl carrier protein (AccB) homologous to MmdC is also expressed by *T. forsythia*, so we checked for the exposure of AccB in intact and sonicated *T. forsythia* cells using a streptavidin-HRP conjugate that recognizes biotinylated proteins, with *P. gingivalis* as a positive control (**Fig. 2A, right panel**). A weak but distinct signal from one or more biotinylated proteins was detected in the intact *T. forsythia* cells regardless of the presence of miropin. The signal was strongly amplified in cell lysates, suggesting *T. forsythia* biotinylates other surface proteins in addition to AccB. These findings show that the integrity of *T. forsythia* cells is not disrupted by dot blotting. Finally, the cell surface exposure of miropin was confirmed by flow cytometry with wt-*Tf* and anti-miropin antibodies, whereas no signal was observed with streptavidin-FITC (**Fig. 2B**). These results show that miropin is displayed on the *T. forsythia* surface but this does not necessarily mean the inhibitor is poised to react with target proteases.

**Figure 2.**
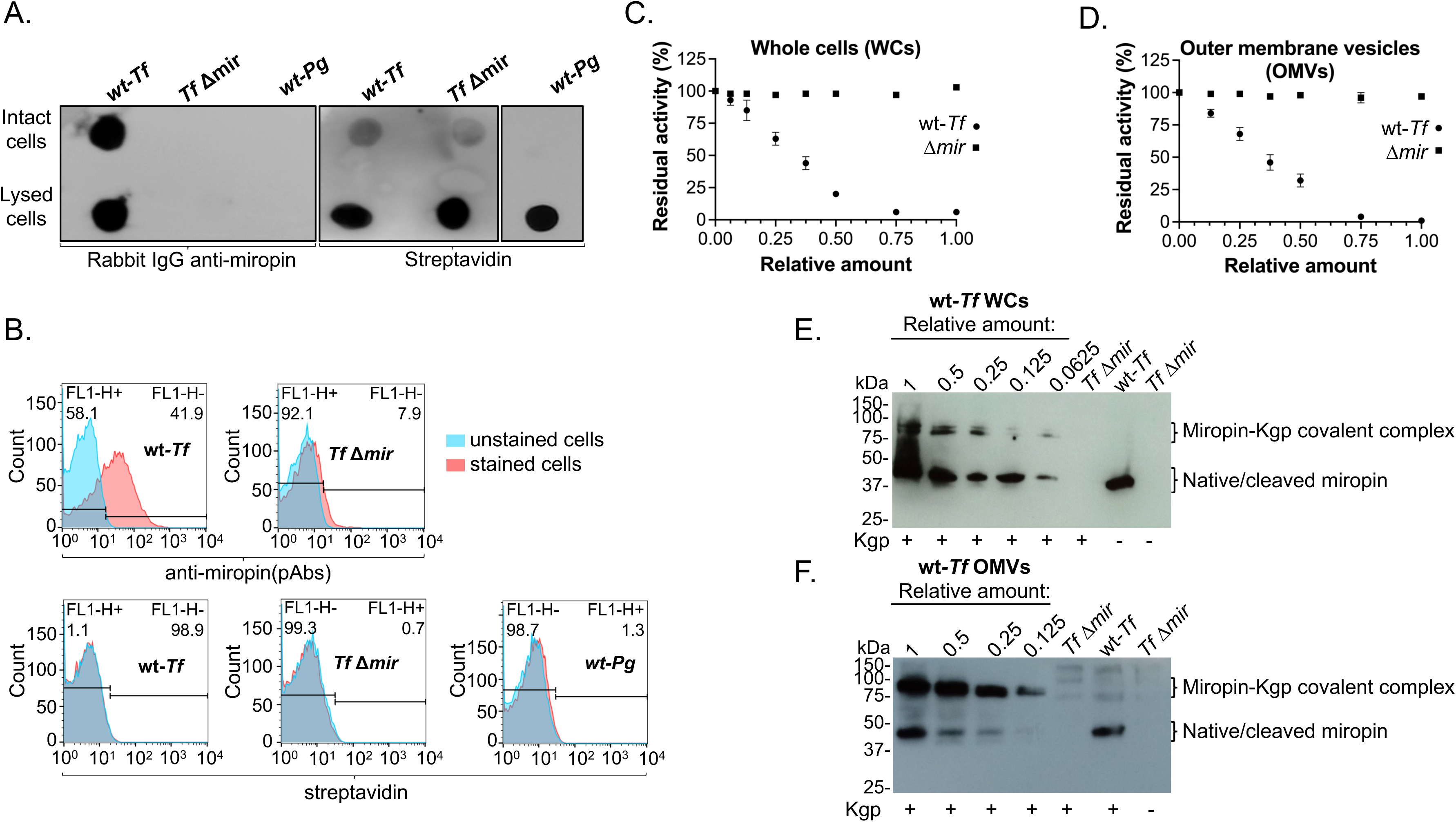
Miropin is located on the *T. forsythia* cell surface and inhibits Kgp. (**A**) Dot blot of intact and sonicated *T. forsythia* cells (wt-*Tf* and *Tf*Δ*mir*) spotted onto nitrocellulose membranes and probed with anti-miropin antibodies followed by HRP-conjugated IgG-specific secondary antibodies (left panel) or streptavidin-HRP (right panels). *P. gingivalis* W83 (wt-*Pg*) as used as a control. (**B**) Flow cytometry for the analysis of wt-*Tf* and *Tf*Δ*mir* stained with either anti-miropin antibodies followed Alexa Fluor 488-conjugated IgG-specific secondary antibodies (upper panels) or streptavidin-FITC (lower panels). Isotype-negative controls are shown in blue. The histograms are representative of three independent experiments. (**C**,**D**) *Kgp* was pre-incubated with increasing amounts of (**C**) wt-*Tf* and *Tf*Δ*mir* cell suspension adjusted to OD_600_ = 1.0 or (**D**) OMV (4.5 mg/mL) derived from the same strains. The residual activity of Kgp was determined and plotted against the volume (μL) of *T. forsythia* cells and OMV suspension. The quantitative data are means ± SEM (n = 3). To visualize covalent inhibitory complex formation, Kgp was mixed with increasing amounts of whole cells (**E**) or OMV (**F**) followed by immumoblot analysis with anti-miropin antibodies followed by HRP-conjugated IgG-specific secondary antibodies (**D**).

The *T. forsythia* surface is covered with a semicrystalline S-layer (Posch *et al*., 2011), which may sterically interfere with protease access to the RCL of miropin. We therefore tested whether Kgp can be inhibited by suspensions of intact *T. forsythia* cells and OMVs (a surrogate for the outer membrane of diderm bacteria). A constant amount of Kgp was pre-incubated with an increasing volume of whole bacterial cells or OMV suspension, and we measured residual Kgp activity. Both cells and OMVs inhibited Kgp in a concentration-dependent manner (**Fig. 2C,D**). This finding was supported by immunoblotting experiments showing the formation of fewer covalent complexes when the volume of cell suspension (**Fig. 2E**) or OMVs (**Fig. 2F**) was reduced. No inhibition or complex formation was observed with the *Tf*Δ*mir* mutant strain or OMVs derived from it. The linear correlation between Kgp activity and the number of *T. forsythia* cells or the protein content of OMV suspensions allowed us to estimate that each *T. forsythia* cell displays 740–1480 miropin molecules, and that OMVs contain 61–116 fmol (1.3–2.5 fg) of miropin per microgram of protein. These data show that a large amount of miropin is present on the *T. forsythia* cell surface and can fully inhibit Kgp activity via the formation of irreversible covalent complexes. This further suggests that miropin plays a role in the reciprocal interactions between *P. gingivalis* and *T. forsythia,* possibly affecting *P. gingivalis* fitness and growth.

### Inhibition of gingipain activity by miropin attenuates the *P. gingivalis* growth *in vitro*

*Porphyromonas gingivalis* is a fastidious, asaccharolytic organism that uses peptides as its source of carbon and energy, and heme is required as a growth factor (Moradali and Davey, 2021). Kgp is important for heme acquisition (Smalley *et al.*, 2011), and gingipains collectively are needed to generate peptide nutrients that *P. gingivalis* takes up via the transporter RagAB (Madej *et al*., 2020). For this reason, the gingipain-null mutant strain of *P. gingivalis* cannot grow on defined media with albumin as the sole source of peptides. We rationalized that miropin, by inhibiting gingipain (Kgp) activity, would limit the growth of *P. gingivalis* on albumin. Accordingly, we found that wild-type VKT-miropin partially but significantly limited wt-*Pg* growth in a concentration-dependent manner. The lower growth rate was comparable to that of the *Pg*Δ*kgp* strain (**Fig. 3A and 3C**).

**Figure 3.**
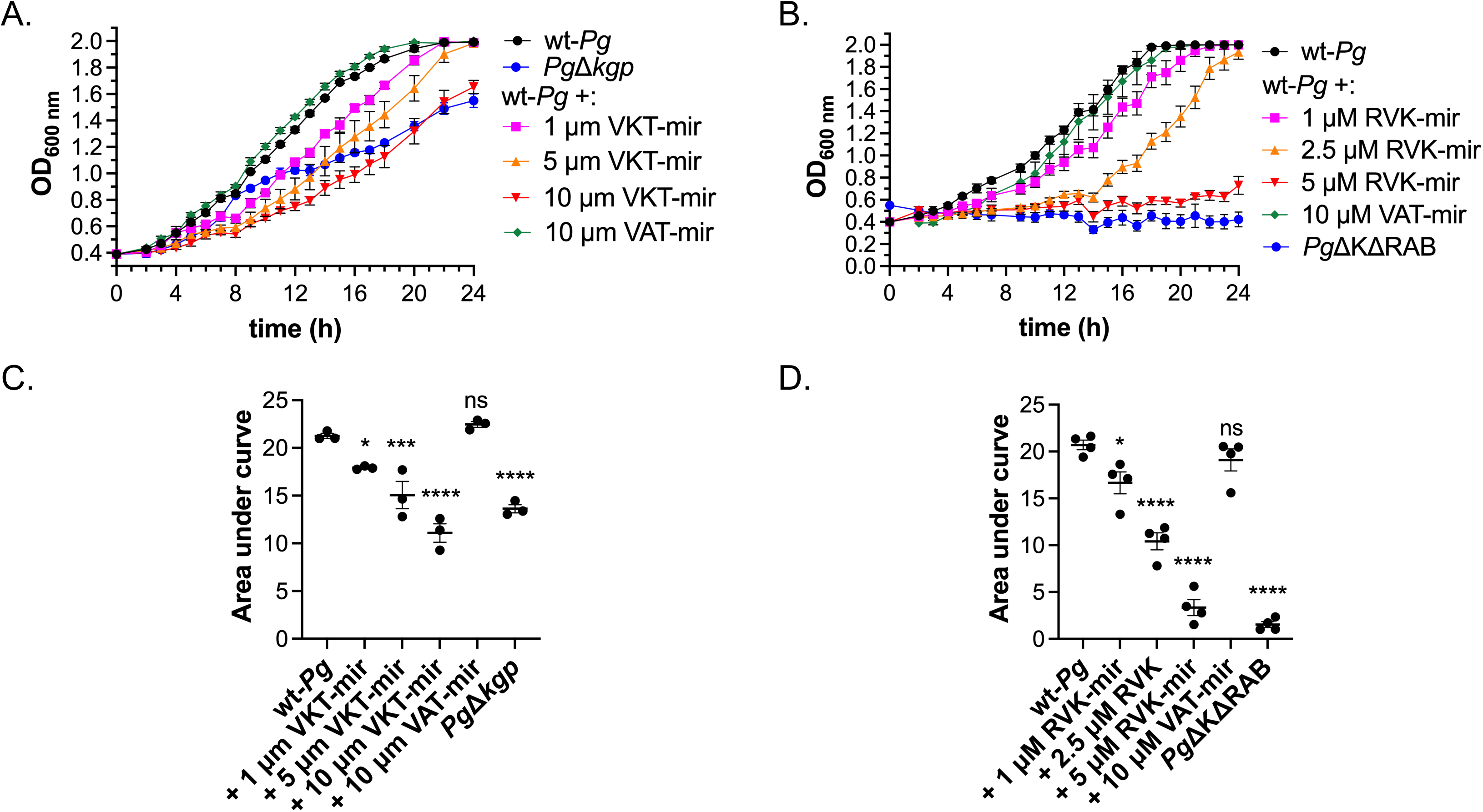
Gingipain-inhibiting variants of miropin slow down the growth of *P. gingivalis* in minimal medium. Growth of *P. gingivalis* W83 (wt-*Pg*), alone or in the presence of different concentrations of recombinant wild-type miropin (**A**) and RVK-miropin (RVK-mir) (**B**) in minimal medium comprising DMEM, 1% BSA, hemin, L-cysteine and menadione was monitored for up to 24 h. To control for the dependence of *P. gingivalis* growth on gingipain activity, wt-*Pg* was cultivated in the presence of VAT-miropin and mutant strains deficient for Kgp (*Pg*ΔKgp) (**A**) and all three gingipains (*Pg*ΔKΔRAB) (**B**). Data are means ± SEM (n = 3).

The inability of wild-type miropin (VKT-miropin) to completely arrest *P. gingivalis* growth may reflect its inability to inhibit arginine-specific gingipains (Rgps = RgpA and RgpB) in keeping with the absence of an Arg residue in the RCL. We therefore exploited the plasticity of miropin for substitutions in the RCL (Goulas *et al*., 2017) to develop miropin variants that inhibit gingipain activity completely. RVK-miropin inhibited both Kgp and RgpB with an SI of 1.7 in both cases (Suppl. Fig. **S2A,B**) and *k_ass_* values of 3.3 x 10^5^ and 7.4 x 10^5^ M^-1^ s^-1^, respectively (Suppl. **Fig. S2C,D**). Conversely, VAT-miropin (with Ala replacing Lys at the RSB) showed no activity against gingipains (not shown) but was a potent inhibitor of NE (Suppl. **Fig. S2E**). RVK-miropin inhibited *P. gingivalis* growth in a concentration-dependent manner, and achieved complete inhibition at 5 µM (**Fig. 3B,D**), resulting is a growth rate similar to that of the gingipain-null mutant strain (*Pg*ΔKΔRAB). In contrast, VAT-miropin had no effect on *P. gingivalis* growth even at 10 µM (**Fig. 3**). To confirm the specificity of inhibition, we measured residual gingipain activity in cultures after 24 h and observed the complete and stable inhibition of Kgp even at the lowest concentration of VKT-miropin and RVK-miropin. In contrast, Rgp activity declined gradually in the presence of increasing concentrations of RVK-miropin, but not VKT-miropin (Suppl. **Fig. S3**). The effect of miropin on *P. gingivalis* can therefore be attributed to the inhibition of gingipains rather than non-target effects resulting from the presence of the miropin protein in the media. This suggests miropin could influence the fitness or virulence of *P. gingivalis in vivo*, because this periodontal pathogen relies on the acquisition of heme and peptides via the proteolytic activity of Kgp (Lewis *et al*., 1999; Smalley *et al*., 2006, 2007, 2008, 2011).

### Recombinant miropin attenuates *P. gingivalis* virulence *in vivo*

To determine whether the inhibition of *P. gingivalis* growth by miropin is relevant *in vivo*, we used the subcutaneous chamber model to examine how our recombinant miropin variants affected *P. gingivalis* proliferation and pathogenicity in the context of a host antimicrobial immune response (O’Brien-Simpson et al, 2003; Maresz *et al.*, 2013). The subcutaneous implantation of titanium coils in this model generates a fibrous chamber with a hollow interior. Introducing bacteria or other mediators into these chambers elicits an inflammatory response and immune cell recruitment in an attempt to clear the bacteria (Maresz *et al.*, 2013).

As shown in **Fig. 4A**, a lethal dose of 1 x 10^9^ CFU *P. gingivalis* W83 introduced into the chamber killed all the animals within 24 h. However, co-injecting a single dose of the wild-type VKT-miropin partially protected the mice, enabling 80% of the animals to survive for 48 h. Surprisingly, and contrary to our observations *in vitro* (**Fig. 3**), wild-type VKT miropin did not reduce the CFU of *P. gingivalis* in the chambers, suggesting it had no effect on proliferation (**Fig. 4B**). This short-lived protective effect possibly reflected the inhibition of Kgp’s pathogenic potential by miropin, as evidenced by the much lower Kgp activity in the chamber exudate fluid (**Fig. 4C**).

**Figure 4.**
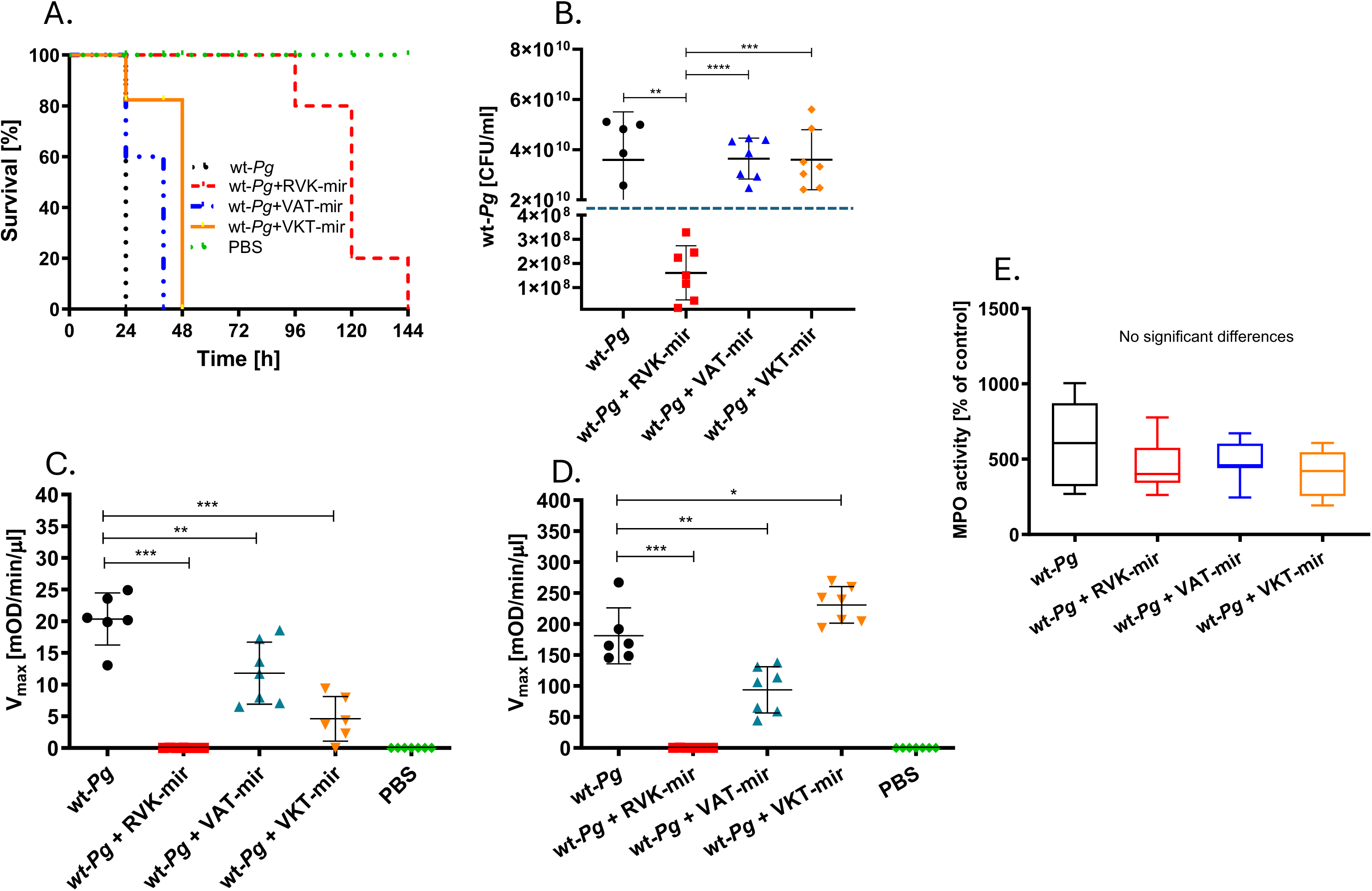
Recombinant miropin protects mice from mortality caused by infection with *P. gingivalis* in an inhibitor-specific manner. Subcutaneous chambers were inoculated with 1.0 × 10^9^ CFU of wild-type *P. gingivalis* (wt-*Pg*) either alone or with added VKT-mir, RVK-mir and VAT-mir. (**A**) Kaplan–Meier survival curves of mice for 6 days post-infection. (**B**-**E**) Chamber fluid (20 µL) was collected 24 h post-infection and analyzed for (**B**) *P. gingivalis* living cells (CFU), (**C**) Kgp, (**D**) Rgp, and (**E**) MPO activity. Each data point represents one mouse, and data from 5–8 mice per group are shown as means ± SD. A vertical dashed line indicates 1 × 10^9^ CFU of *P. gingivalis* in the inoculum at T_0_. Comparative analysis revealed that bacterial counts changed significantly (p < 0.001) at 24 h in all cases compared to the initial inoculum. Statistically significant differences in bacterial load were observed between wt-*Pg* and wt-*Pg* + RVK-mir (**p < 0.01), between wt-*Pg* + RVK-mir and wt-*Pg* + VAT-mir (****p < 0.0001), and between wt-*Pg* + RVK-mir and wt-*P*g + VKT-mir (***p < 0.001). Statistical significance was determined by Welch’s t-test or one-way ANOVA followed by Dunnett’s multiple comparisons test.

In contrast to wild-type VKT-miropin, RVK-miropin enabled the mice to survive for 96 h (**Fig. 4A**), and significantly reduced the *P. gingivalis* CFU in the chamber fluid (**Fig. 4B**). This reflected the inhibition of both Kgp and Rgp (**Fig. 4C,D**), which affected both the survival and growth of *P. gingivalis* in the chambers. As expected, VAT-miropin offered no significant protection against *P. gingivalis* (**Fig. 4A**), correlating with a higher *P. gingivalis* CFU after 24 h (**Fig. 4B**). Interestingly, despite its inability to inhibit gingipains, VAT-miropin nevertheless reduced the activity of Rgp (**Fig. 4C**) and Kgp (**Fig. 4D**) in the chambers. We also measured the levels of MPO as a marker of neutrophils infiltrating into the chambers. MPO levels increased following *P .gingivalis* infection, but MPO activity was reduced in chambers injected with miropin regardless of the variant, but this reduction was not significant (**Fig. 4E**). These results once again confirm that gingipains are key *P. gingivalis* virulence factors contributing to its pathogenicity in the chamber model of infection (Benedyk *et al*., 2019).

### Co-infection with *T. forsythia* abrogates *P. gingivalis* pathogenicity in the chamber model and prevents alveolar bone loss in the oral gavage model

Given that *P. gingivalis* and *T. forsythia* coexist naturally in periodontal pockets, we compared the pathological outcomes of mono-infection versus co-infection with different strains, and assessed the impact of miropins on their combined virulence using the chamber model. We hypothesized that miropin displayed by *T. forsythia* cells would inhibit Kgp and Tpr on the surface of *P. gingivalis*, thereby reducing the pathogenic potential of *P. gingivalis in vivo*, which is mainly dependent on Kgp in mouse models of infection (O’Brien-Simpson *et al*., 2001; Pathirana *et al.*, 2007).

To test our hypothesis, we first demonstrated that the activities of Kgp and Tpr were reduced to 5% and 10%, respectively, in a 1:1 suspension of *T. forsythia* and *P. gingivalis*, each adjusted to OD_600_ = 1 (Fig. **S4A**). The formation of covalent miropin-Kgp complexes was confirmed by immunoblotting with anti-miropin antibodies (Suppl. **Fig. S4B**). Mice were then infected by direct injection into subcutaneously implanted chambers with 10^9^ CFU wt-*Pg* or wt-*Tf* alone or a combination of wt-*Pg* and the mutant *TfΔmir*. All mice inoculated with *P. gingivalis* alone died within 24–36 h whereas those infected with wt-*Tf* alone or both wild-type bacteria showed no signs of illness for up to 6 days post-infection (**Fig. 5A**). This protective effect in the co-infection model was attributed to the inhibition of Kgp by miropin because mice co-infected with *P. gingivalis* and *TfΔmir* became ill and died at the same rate as those infected solely with *P. gingivalis* (**Fig. 5A**). The *T. forsythia* strains had no significant effect on *P. gingivalis* growth in the chambers (**Fig. 5B**), correlating with similar levels of Rgp activity in the chamber fluid (**Fig. 5C**) despite Kgp activity falling to baseline in chambers inoculated with both wild-type bacteria (**Fig. 5D**). The latter confirms that Kgp activity is inhibited by miropin on the surface of *T. forsythia* even *in vivo*. The only other notable effect of co-infection compared to mono-infection with *P. gingivalis* was the reduced number of infiltrating neutrophils, as indicated by MPO levels (**Fig. 5E**).

**Figure 5.**
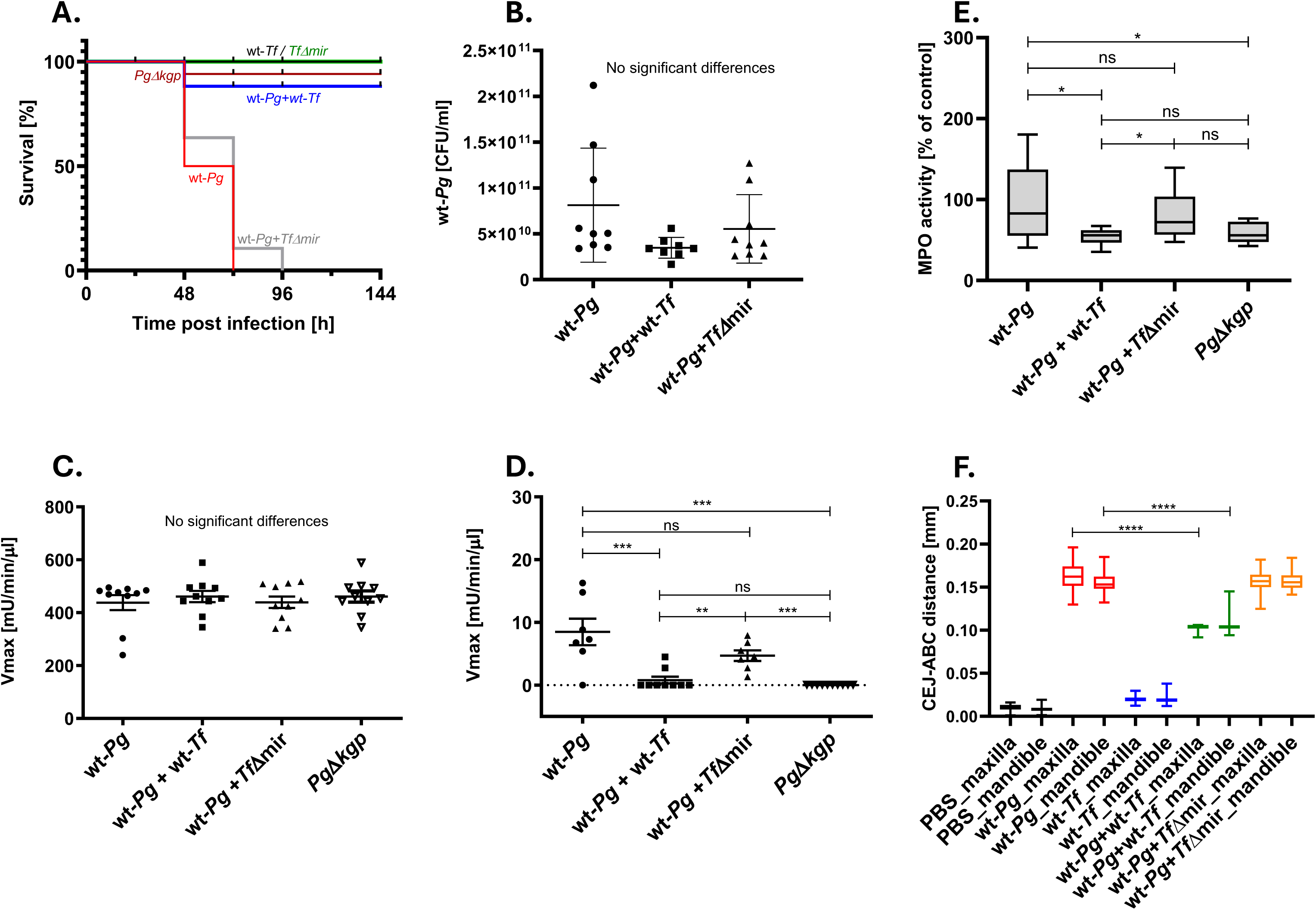
Co-infection with *T. forsythia* abrogates *P. gingivalis* pathogenicity in the chamber model and prevents alveolar bone loss in the oral gavage model. (**A-E**) Subcutaneous chambers were inoculated with *P. gingivalis* W83 (wt-*Pg*), wild-type *T. forsythia* (wt-*Tf*) and miropin-null strains (*Tf*Δmir), or Kgp-deficient *P. gingivalis* W83 (*Pg*Δkgp) alone at 1 x 10^9^ CFU, or in the combinations wt-*Pg* + wt-*Tf* and wt-*Pg* + *Tf*Δmir. Mortality of infected mice was recorded for 6 days (**A**). We sampled 20 μL of chamber fluid 24 h post-infection and determined the number of living *P. gingivalis* cells by plating and colony counting (**B**), as well as measuring Rgp (**C**) and Kgp (**D**) activities. MPO activity was determined as a surrogate of neutrophil infiltration (**E**). (**F**) Mice were infected orally with wt-*Pg* alone (1 x 10^10^ CFU) or in combination with wt-*Tf* or *Tf*Δ*mir*, each at 1 x 10^10^ CFU. Control mice were infected orally with wt-*Tf* or *Tf*Δmir. Alveolar bone loss was determined by using micro-computed tomography (µCT) to measure the distance from the cementoenamel junction (CEJ) to the alveolar bone crest (ABC). Data are presented as mean ± SD, except for Fig. 5C and Fig. 5D, where values are shown as mean ± SEM (n = 8-10). Statistical significance was determined using Welch’s *t*-test and one-way ANOVA followed by Dunnett’s multiple comparisons test (*p<0.05, **p<0.01, ***p<0.001, ns = not significant).

In periodontitis lesions, *T. forsythia* and *P. gingivalis* exist in close proximity and both are tightly associated as aggregates inside cells of the stroma, squamous epithelium, and capillary endothelium (Rajakaruna *et al.*, 2018). This close relationship suggests that the inevitable inhibition of Kgp by miropin will limit the ability of *P. gingivalis* to cause alveolar bone loss during co-infection with *T. forsythia* in the murine periodontitis model. To test this hypothesis, we infected mice orally with *P. gingivalis* alone or combined with wt-*Tf* or *Tf*Δ*mir*. Control mice were also infected orally with wt-*Tf or Tf*Δ*mir*. Neither of the *T. forsythia* strains caused any significant bone loss, but *P. gingivalis* infection resulted in the measurable erosion of the alveolar bone (**Fig. 5F**). Interestingly, co-infection with *T. forsythia* significantly reduced *P. gingivalis*-induced bone loss, but the effect was dependent on miropin expression because the *Tf*Δ*mir* strain offered no protection. We found that *T. forsythia* was not pathogenic in the oral gavage model, where alveolar bone loss is the main indicator. But miropin on the *T. forsythia* surface or released in soluble form was able to attenuate *P. gingivalis* virulence during co-infection, consistent with the important role of Kgp in bone loss caused by *P. gingivalis* in this murine periodontitis model (Pathirana *et al.*, 2007).

## DISCUSSION

Our understanding of the microbial causes of periodontitis has evolved from a non-specific bacterial challenge involving the red complex pathogens (*P. gingivalis*, *T. forsythia* and *Treponema denticola*) to a concept of microbial dysbiosis, where synergistic interactions within a broad community of microbes, including commensals and pathobionts, play a key role (Han and Ding 2025). In this newer paradigm, periodontitis is initiated in susceptible hosts by the growth of keystone pathobionts, which corrupt the otherwise commensal microbes in the subgingival biofilm to create a dysbiotic microbiome (Hajishengallis *et al.*, 2023). The host immune response cannot disassemble this dysbiotic community, triggering a feed-forward loop of inflammation, resulting in the erosion of periodontal tissues. Proteolytic enzymes secreted by periodontal pathogens, especially gingipains such as Kgp, play a key role in this process. Discovering that Kgp is inhibited by miropin provides new insights into the combined pathogenic potential of *P. gingivalis* and *T. forsythia* within the dysbiotic biofilm.

Both pathogens are thought to aggregate in the subgingival biofilm, where they show evidence of metabolic interdependence (Shimotahira *et al*., 2013; Zhu and Lee, 2016). For example, *T. forsythia* extracts promote the growth of *P. gingivalis* (Yoneda *et al*., 2005; Ng *et al*., 2016), whereas *T. forsythia* is an *N*-acetylmuramic acid auxotroph that requires scavenged external bacterial components to support its peptidoglycan synthesis and growth (Mayer et al., 2019; Wodzanowski et al., 2022). The role of gingipains in this interspecies interaction is evident from two key observations. First, the abundance and distribution of *T. forsythia* in experimental multispecies biofilms are influenced by the secretion of Kgp by *P. gingivalis* (Bao *et al*., 2014). Second, efficient co-invasion of gingival keratinocytes by these species depends on gingipain activity (Jung *et al*., 2016). Overall, these findings suggest that miropin may influence the virulence of *P. gingivalis* .

Miropin targets proteases with widely different specificities by using three distinct peptide bonds at the RCL: Val^367^-Lys^368^, Lys^368^-Thr^369^ and Thr^369^-Ser^370^ (Goulas *et al*., 2017). A similar mechanism has been reported for α_2_-antiplasmin, which inhibits plasmin and chymotrypsin at Arg^364^-Met^365^ and Met^365^-Ser^366^, respectively (Potempa *et al*., 1988). However, none of the known natural serpins uses three peptide bonds to expand their range of target proteases. Most serpins inhibit serine proteases, but some, such as viral serpin (cytokine response modifier A) and the human squamous cell carcinoma antigens SCCA1 (SERPINB3) and SCCA2 (SERPINB4), inhibit different members of the cysteine proteinase class with a papain-like fold (Schick *et al*., 1998; Luke *et al*., 2000). Even so, miropin is unique in inhibiting serine proteases from the trypsin and subtilisin families, as well as papain-like, caspase-like (gingipains), and calpain-like (Tpr) cysteine proteases. It is tempting to speculate that these unique features of miropin evolved to protect *T. forsythia* surface proteins from cleavage by host and bacterial proteases in the crowded microbial environment of inflamed periodontal pockets.

Miropin is strongly expressed *in vivo* in *T. forsythia*-infected periodontal pockets (Eckerd *et al.*, 2018). We determined the abundance of this inhibitor on the cell surface (740–1480 molecules per cell), where it is anchored to the outer membrane as a lipoprotein. This positioning allows it to inhibit not only soluble Kgp but also gingipains on the surface of *P. gingivalis*. The inhibition of Kgp is kinetically efficient, allowing VKT-miropin to slow the growth of *P. gingivalis* on defined media *in vitro* with albumin as the only nutrient source, in a concentration-dependent manner. VKT-miropin also significantly reduced Kgp activity *in vivo* in the chamber infection model, extending mouse survival by 24 h. However, residual activity appeared sufficient to support the growth of *P. gingivalis* growth over the longer term, ultimately with lethal effects.

Compared to VKT-miropin, RVK-miropin (which inhibits both Kgp and Rgp) provided much greater protection, reducing *P. gingivalis* growth in the chamber and extending the survival of infected mice to 4–5 days. The data suggest Rgps contribute significantly to *P. gingivalis* pathogenicity in this infection model. Alternatively, inhibiting both gingipains with RVK-miropin may cause bacterial starvation, as evidenced by the complete inhibition of *P. gingivalis* growth *in vitro* when albumin is the only nutrient source. As a control, VAT-miropin had a minimal effect on mouse survival and did not impede *P. gingivalis* proliferation in the chamber.

The claim that *T. forsythia* can restrict *P. gingivalis* virulence was confirmed using the chamber and oral gavage models. In both cases, mice co-infected with wt-*Tf* and wt-*Pg* survived for longer and lost significantly less alveolar bone mass compared to mice infected with wt-*Pg* alone. The same levels of survival and bone preservation were observed in mono-infections with *Pg*Δ*kgp*. However, co-infection with *Tf*Δ*mir* did not affect *P. gingivalis*-induced pathology. In the chamber model, wt-*Tf* did not affect *P. gingivalis* proliferation or Rgp activity but did reduce Kgp activity to background levels. Interestingly, neutrophil infiltration was significantly reduced in chambers inoculated with wt-*Tf* and wt-*Pg*, reaching the levels seen with *Pg*Δ*kgp*, compared to infection with *P. gingivalis* alone or co-infection with *Tf*Δ*mir*. This correlated with mouse survival, arguing that Kgp activity plays a key role in lethal inflammation in the chamber model and confirming Kgp as a leading factor contributing to alveolar bone loss in experimental periodontitis (Pathirana *et al*. 2007), and morbidity in the murine lesion model (O’Brien-Simpson *et al*., 2000).

The three species *P. gingivalis*, *T. denticola* and *T. forsythia* are recognized as key periodontal pathogens due to their frequent co-localization in deep periodontal pockets (Ximénez-Fyvie et al., 2000ab). However, the specific role of *T. forsythia* in the initiation and progression of periodontitis remains unclear (Yang *et al.*, 2004; Moradi *et al.*, 2025). The pathogenicity of this species has been linked to the expression of virulence factors such as the leucine-rich repeat protein BspA, the apoptosis-inducing cysteine protease PrtH, surface-layer glycoproteins (TfsA, TfsB), glycosidases (sialidases and neuraminidases) (Sharma, 2010), and the KLIKK family of proteases (Ksiazek *et al*., 2015a). These molecules are thought to target specific components of the host immune system, helping *T. forsythia* to survive and promoting inflammatory processes.

Thus far, only the cell surface-associated protein BspA has been directly linked to alveolar bone loss in a murine model of periodontitis (Sharma *et al.*, 2005). This effect was more prominent in a co-infection model, where *T. forsythia* was combined with other periodontal pathogens such as *Fusobacterium nucleatum* (Settem *et al.*, 2012). In rats, oral co-infection with *T. forsythia*, *P. gingivalis* and *T. denticola*, with or without *F. nucleatum*, significantly increased alveolar bone loss compared to mono-infections (Kesavalu *et al.*, 2007). Synergistic effects have also been observed for *T. forsythia* and *P. gingivalis* in skin abscess models in rabbits (Takemoto *et al*., 1997) and mice (Yoneda *et al*., 2001), as well as additive effects in a rat model of periodontitis (Verma *et al*., 2010). In contrast, we observed no pathological effects caused by *T. forsythia* mono-infection in two separate *in vivo* models. This discrepancy may reflect differences in the strains of *T. forsythia* used, the type of infection model (e.g., skin abscess vs chamber model), or variations in bacterial preparation and inoculum size.

Despite the above disparity, there is little doubt that secreted miropin has the potential to reduce excessive proteolytic activity caused by host proteases in the inflamed periodontal pocket (Kennett *et al.*, 1997). Miropin is a highly effective inhibitor of neutrophil serine proteases, especially NE and CatG (Książek *et al.*, 2015). Elevated NE activity is strongly linked to deep periodontal pockets (Figueredo and Gustafsson, 1998) and associated with the presence of subgingival periodontal pathogens (Jin et al., 1999; Gul et al., 2016). Elastase activity in gingival crevicular fluid (GCF) acts as a predictor of future attachment loss (Eley and Cox, 1996) and serves as a biomarker for periodontal treatment success (Gul *et al*., 2017). Elastase is a highly aggressive protease than can contribute to the development and progression of periodontal disease through multiple mechanisms (Alfakry *et al*., 2016; Tseng *et al.*, 2022). Therefore, its inhibition by miropin may offer protective benefits. Conversely, protecting *T. forsythia* cell-surface-associated virulence factors from proteolytic degradation could enhance the survival and virulence of this pathogen, especially within the dysbiotic biofilm immersed in the highly proteolytic environment of GCF.

In conclusion, we cannot unambiguously exonerate *T. forsythia* from active involvement in the pathobiology of periodontitis, but following the legal principle of “innocent until proven guilty,” one could argue that *T. forsythia* is simply an innocent bystander coexisting with true periodontal pathogens.

## ACKNOWLEDGEMENTS

This study was supported by NIDCR grant DE030939 to JB and JP, and National Science Centre (NCN, Poland) grant 2015/17/B/NZ1/00666 to MK.

