## Supplemental Figures for "Innocent until proven guilty: *Tannerella forsythia* may attenuate the virulence of *Porphyromonas gingivalis*"

Suppl. Fig. S1

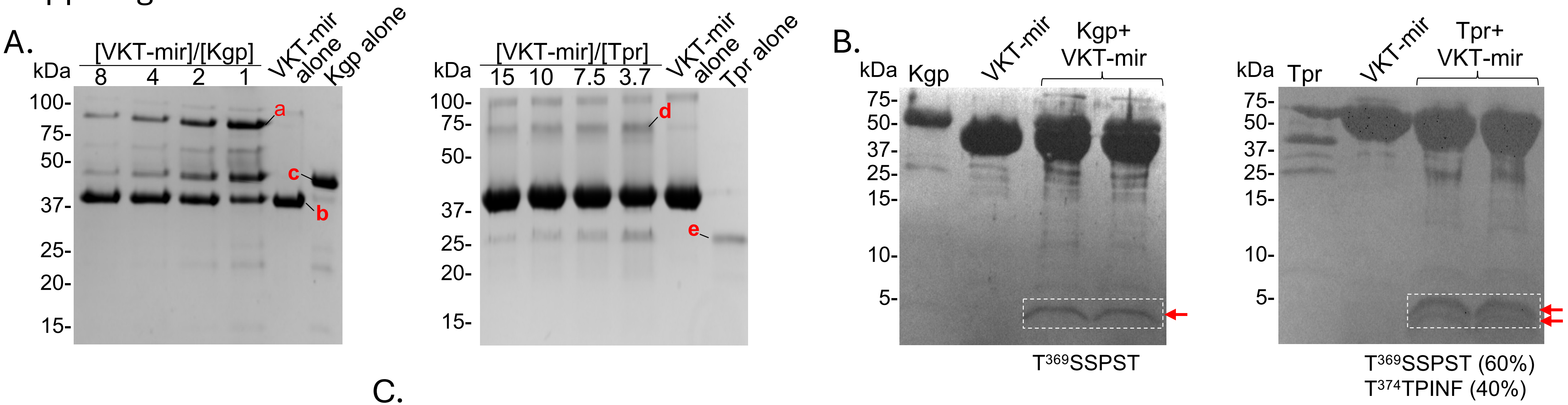


C.

| Band | Protein ID by MS-MS | Protein score |
|---|---|---|
| 1 | miropin (*T. forsythia*)<br>Kgp (*P. gingivalis*) | 2392<br>1037 |
| 2 | miropin (*T. forsythia*) | 7464 |
| 3 | Kgp (*P. gingivalis*) | 2892 |
| 4 | miropin (*T. forsythia*)<br>Tpr (*P. gingivalis*) | 2036<br>492 |
| 5 | Tpr (*P. gingivalis*) | 353 |

D.

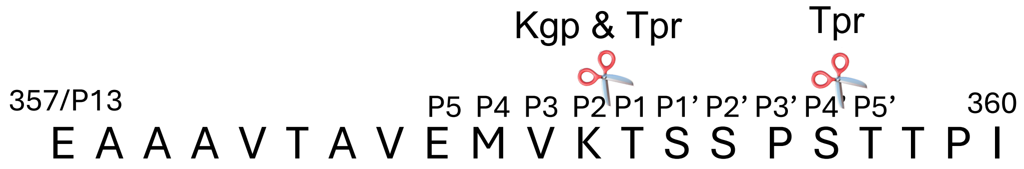


**Figure S1. Miropin forms covalent complexes with *P. gingivalis* proteases Kgp and Tpr.** Miropin was incubated for 15 min with proteases at different molar ratios (**A**) or at the 1:1 ratio (**B**) and resolved by SDS-PAGE using Laemmli and Tris-Tricine buffer systems, respectively. (**B**) Bands of high molecular mass complexes (a and d), miropin (b), Kgp (c), and Tpr (e) were excised from gels and analyzed by peptide mass fingerprinting. The best total MS/MS scores are presented (**C**). A ~5 kDa peptides (arrows on panel **B**) were cut from the gels and subjected to N-terminal sequence analysis. Identified cleavage sites at the RCL are shown (**D**).

Suppl. Fig. 2

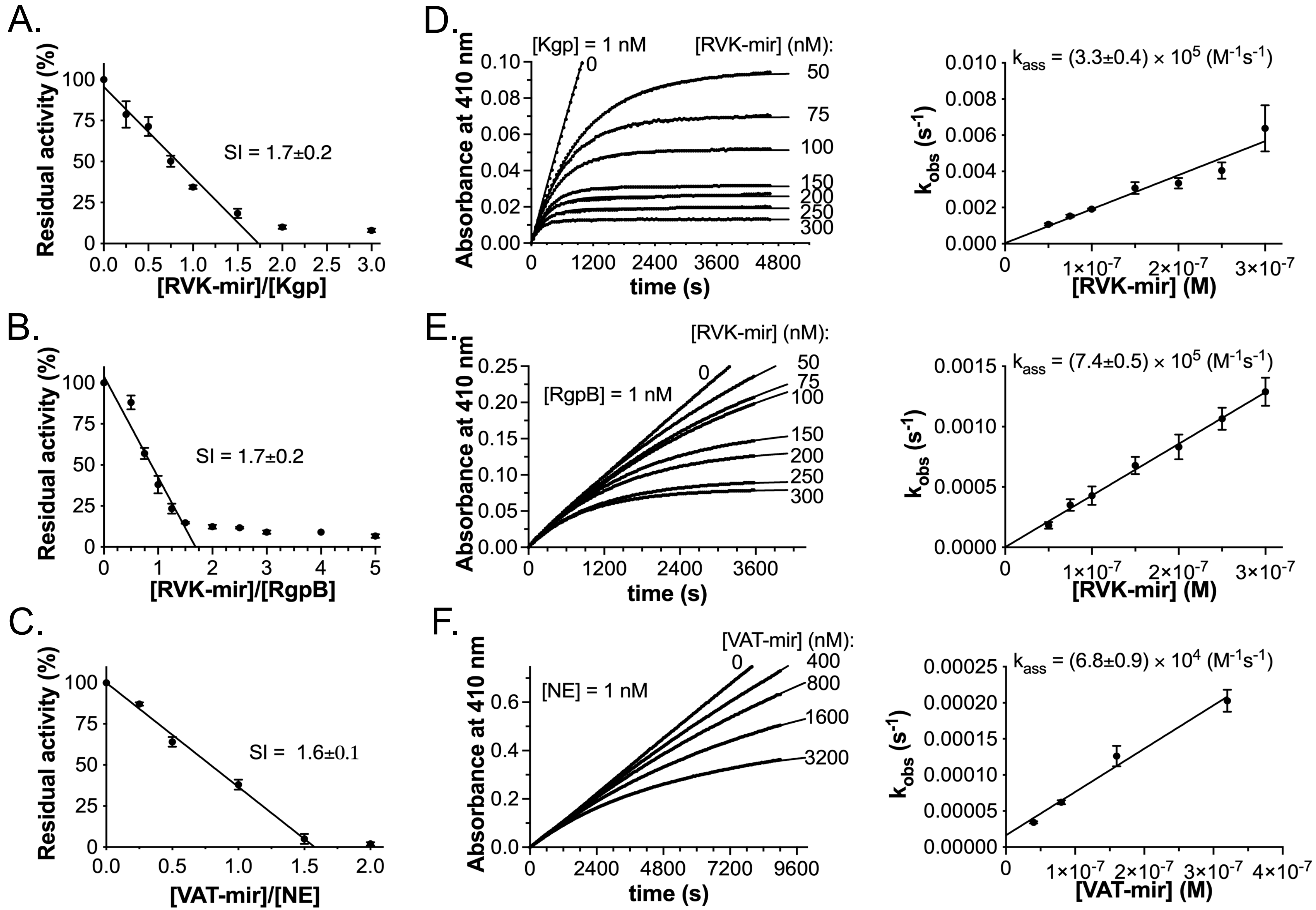

A.
Residual activity (%)
SI = 1.7±0.2
[RVK-mir]/[Kgp]
B.
Residual activity (%)
SI = 1.7±0.2
[RVK-mir]/[RgpB]
C.
Residual activity (%)
SI = 1.6±0.1
[VAT-mir]/[NE]
D.
[Kgp] = 1 nM
[RVK-mir] (nM):
Absorbance at 410 nm
time (s)
$k_{ass}$ = (3.3±0.4) × $10^5$ ($M^{-1}s^{-1}$)
$k_{obs}$ ($s^{-1}$)
[RVK-mir] (M)
E.
[RVK-mir] (nM):
[RgpB] = 1 nM
Absorbance at 410 nm
time (s)
$k_{ass}$ = (7.4±0.5) × $10^5$ ($M^{-1}s^{-1}$)
$k_{obs}$ ($s^{-1}$)
[RVK-mir] (M)
F.
[VAT-mir] (nM):
[NE] = 1 nM
Absorbance at 410 nm
time (s)
$k_{ass}$ = (6.8±0.9) × $10^4$ ($M^{-1}s^{-1}$)
$k_{obs}$ ($s^{-1}$)
[VAT-mir] (M)

**Figure S2. Substitution of amino acid residues in the VKT sequence in the reactive center loop (RCL) of native miropin changes the spectrum of targeted proteases.** Kgp (**A**), RgpB (**B**), and neutrophil elastase (NE) (**C**) were preincubated with increasing concentrations of RVK-miropin (for Kgp and RgpB) (**A** and **B**) and VAT-miropin for NE (**C**) at 37 °C for 15 min. The residual activity was determined using *N*-tosylo-GPK-pNA for Kgp, Bz-Arg-pNA for RgpB, and Suc-AAPV-pNA for NE, respectively, and plotted against the miropin:protease molar ratio. Activity of enzymes in the absence of inhibitor was considered as 100%. Of note, VAT miropin did not affect Kgp and RgpB activity (not shown). **D-F,** Progress curve analysis was used to determine the $k_{ass}$ values of Kgp (**D**) and RgpB (**E**), and NE (**F**) inhibition by RVK-mir (**D**,**E**) or VAT-mir (**F**). Proteases were added to mixtures containing a constant amount of substrate and increasing concentrations of miropin. Changes in absorbance were then recorded. The number associated with each progress curve represents the concentration of miropin (nM). The values of $k_{obs}$ were plotted as a function of miropin concentration; $k_{ass}$ was determined from the slope of the fitted linear curve to the data points and corrected for the stoichiometry factor. The data presented are mean value ± SD (N = 3).

Suppl. Fig. 3

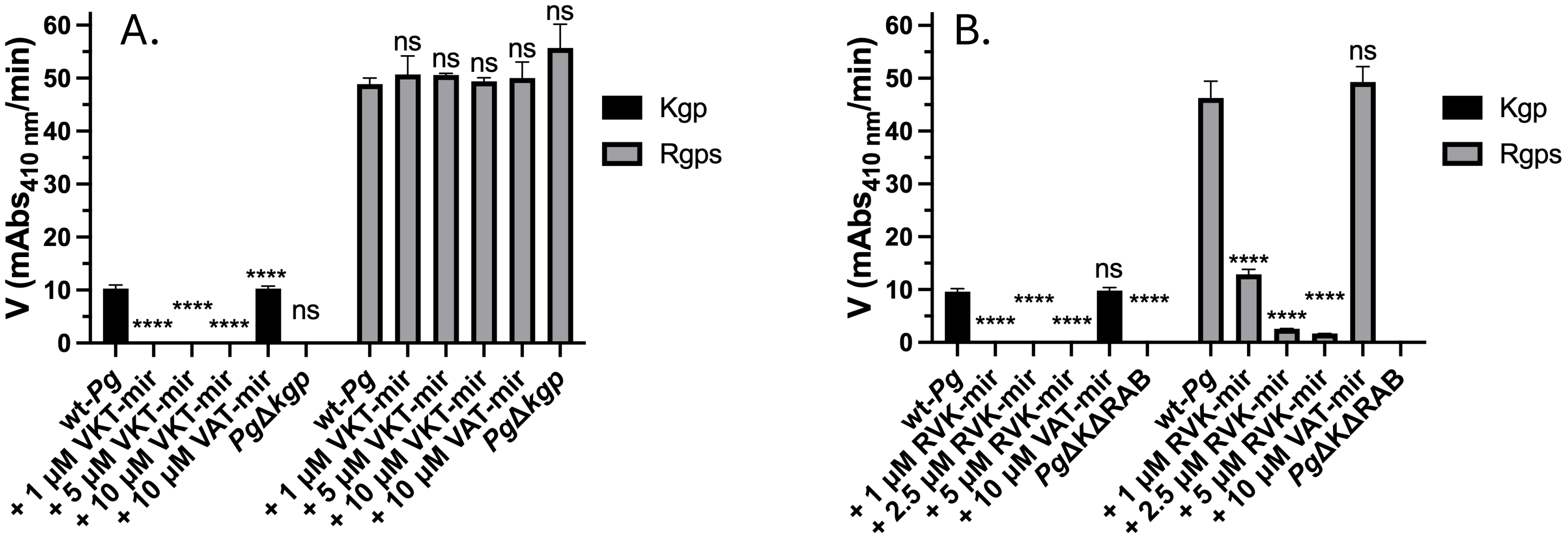


**Figure S3. Gingipain activity when *P. gingivalis* is grown in the presence of miropin.** We cultured *P. gingivalis* W83 (wt-*Pg*) alone or in the presence of different concentrations of recombinant wt-miropin (VKT-mir) (**A** and **B**), RVK-miropin (RVK-mir) (**B**) and VAT-miropin (VAT-mir) (**A** and **B**) under anaerobic conditions in minimal medium comprising DMEM, 1% BSA, hemin, L-cysteine, and menadione for 24 h, and the activity of Kgp and Rgps was measured using Ac-Lys-pNA and BAPNA as substrates, respectively. PgΔ*kgp* and PgΔKΔRAB were used as controls. Data are means ± SEM (n = 3). Group comparisons were performed using one-way ANOVA followed by Dunnett's multiple comparisons test (****p<0.0001 compared with wt-*Pg*, ns, no statistical significance).

Suppl. Fig. S4

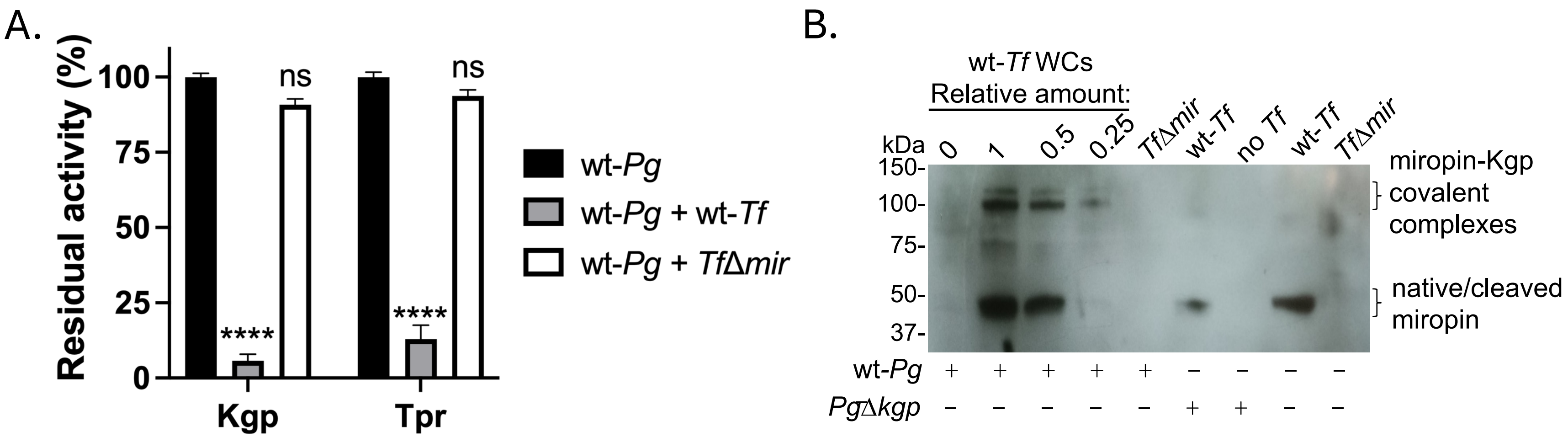


**Figure S4. Intact *T. forsythia* cells inhibit the activity of Kgp and Trp on the *P. gingivalis* cell surface by covalent complex formation in a miropin-dependent manner.** (**A**) Cell suspensions of *P. gingivalis* W83 (wt-*Pg*) and wild-type *T. forsythia* (wt-*Tf*) or its isogenic miropin-null mutant strain (*Tf*Δ*mir*) were adjusted to $OD_{600}$ = 1 and mixed 1:1 for 15 min, then the residual activity of Kgp and Tpr was determined using N-tosylo-GPK-pNA and Suc-LLVY-AMC, respectively, as a substrate. (**B**) Cell suspensions of wt-*Tf* and *Tf*Δ*mir* were mixed with wt-*Pg* or *Pg*Δkgp at the indicated ratio for 15 min, and the reaction was stopped by boiling in non-reducing sample buffer before SDS-PAGE and immunoblotting. The membrane was probed with anti-miropin antibodies followed by HRP-conjugated IgG-specific secondary antibodies, and the blot was developed with ECL Western Blotting Substrate. Samples containing *T. forsythia* cells alone or *P. gingivalis* alone were run as controls.
